# Diversity of free-living prokaryotes on terrestrial and marine Antarctic habitats

**DOI:** 10.1101/2021.04.27.441673

**Authors:** Amanda Gonçalves Bendia, Julio Cezar Fornazier Moreira, Juliana Correa Neiva Ferreira, Renato Gamba Romano, Ivan Gonçalves de Castro Ferreira, Diego Castillo Franco, Heitor Evangelista, Rosalinda Carmela Montone, Vivian Helena Pellizari

## Abstract

Microorganisms in Antarctica are recognized for having crucial roles in ecosystems functioning and biogeochemical cycles. In order to explore the diversity and composition of microbial communities through different terrestrial and marine Antarctic habitats, we analyze 16S rRNA sequence datasets from fumarole and marine sediments, soil, snow and seawater environments. We obtained measures of alpha- and beta-diversities, as well as we have identified the core microbiome and the indicator microbial taxa of a particular habitat. Our results showed a unique microbial community structure according to each habitat, including specific taxa composing each microbiome. Marine sediments harbored the highest microbial diversity among the analyzed habitats. In the fumarole sediments, the core microbiome was composed mainly by thermophiles and hyperthermophilic Archaea, while in the majority of soil samples Archaea was absent. In the seawater samples, the core microbiome was mainly composed by cultured and uncultured orders usually identified on Antarctic pelagic ecosystems. Snow samples exhibited common taxa in comparison to the habitats from the Antarctic Peninsula, which suggests long-distance dispersal processes occurring from the Peninsula to the Continent. This study contributes as a baseline for further efforts on evaluating the microbial responses to environmental conditions and future changes.

## 1. Introduction

Despite extreme conditions, Antarctica harbors a complex mosaic of microbial habitats (Bowman, 2018). In these habitats, microorganisms play a fundamental role in the food web and in the biogeochemical cycles. Recent studies revealed diverse bacterial and archaeal communities inhabiting terrestrial and marine habitats in Antarctica, showing to be distinct from Arctic and alpine communities (Boetius et al., 2015). Terrestrial habitats for free-living prokaryotes in Antarctica include especially mineral, ornithogenic and geothermal soils, permafrost, lakes, glaciers, snow and rocks. The microbial diversity in these habitats have been firstly described using culture-dependent methods (e.g. Friedmann et al., 1988; Hirsch et al., 1988; Siebert et al., 1996; Siebert and Hirsch, 1988), and most recently, through culture-independent strategies, mainly by 16S rRNA sequencing (e.g. Alekseev et al., 2020; Almela et al., 2021; Archer et al., 2019; Bendia et al., 2018; Franco et al., 2017; Malard et al., 2019). These studies have shown phyla such as Proteobacteria, Actinobacteria, Acidobacteria, Bacteroidetes and Firmicutes as abundant in soils and permafrosts from Antarctic Peninsula (Bottos et al., 2014; Jansson and Taş, 2014), whereas Cyanobacteriia, Flavobacteriia and Alphaproteobacteria were the prevalent classes in snow samples from the Antarctic Plateau (Michaud et al., 2014).

Marine habitats generally include deep and shallow sediments, and water column at both euphotic (<200 m) and aphotic zones (>200 m). Signori et al. (2014) studied microbial communities in water column at Bransfield Strait, Southern Ocean, and found Thaumarchaeota, Euryarchaeota and Proteobacteria (Gamma-, Delta-, Beta-, and Alphaproteobacteria) as abundant taxa below 100 m, whereas the dominant phyla above 100 m were Bacteroidetes and Proteobacteria (mainly Alpha- and Gammaproteobacteria). In marine sediments from Admiralty Bay (100–502 m total depth) (King George Island) and adjacent North Bransfield Basin (693–1147 m), Gammaproteobacteria was found as a highly abundant taxa (>90%), followed by Alpha- and Deltaproteobacteria, Firmicutes, Bacteroidetes and Actinobacteria (Franco et al., 2017).

Although previous studies have described microbial communities in different environments from Maritime and Continental Antarctica (e.g. Alekseev et al., 2020; Almela et al., 2021; Archer et al., 2019; Bendia et al., 2018; Cavicchioli, 2015; Cowan et al., 2014; Franco et al., 2017; Malard et al., 2019; Signori et al., 2014), few have focused on indicating the microbiome across a range of Antarctic habitats. In this study, we aimed to reveal the microbiome of five habitats (fumarole sediment, marine sediment, snow, soil and seawater) at two main Antarctic locations, including Antarctic Peninsula (King George Island and Deception Island) and Continental Antarctica (West Antarctica, 670 km from geographical South Pole, near Criosfera 1 module). We were able to describe the core microbiome and the microbial indicators of the different Antarctic habitats, contributing as a baseline study for further efforts on evaluating the microbial responses to environmental conditions and future changes.

## 2. Methodology

### 2.1. Study area and sampling strategy

All the samples selected for this study were collected during the Brazilian Antarctic expeditions (OPERANTAR) XXX to XXXV, comprising the years from 2012 to 2017, and were supported by the following projects: Microsfera (CNPq 407816/2013-5), INCT-Criosfera (CNPq 028306/2009 - Criosfera 1 module) and MonitorAntar (USP-IO/MMA-SBF Agreement No. 009/2012). Detailed information is described in Supplementary Table 1.

The samples selected for this study comprise areas located in both Maritime and Continental Antarctica. In addition, samples include 5 different sample types, comprising the following habitats: marine sediment, fumarole sediment, snow, seawater and soil.

The sampling sites in Maritime Antarctica included King George Island (S 62° 23’ S, W 58° 27’) and Deception Island (S 62° 55’, W 60° 37’), located in the South Shetland archipelago. Samples from King George Island included seawater, marine sediment and soil. Seawater samples were collected at Admiralty Bay near Wanda and Ecology Glaciers, using a Van-Dorn water-sampling bottle. Three water depths were collected and classified as superficial (0 - 5 m), intermediate (~10 m) and bottom (~30 m) depths. Approximately 5 L of water of each sample were filtered on the Brazilian Antarctic Station “Comandante Ferraz” (EACF) using a vacuum pump and 0.22 μm-membrane filters. Superficial marine sediments (0 - 5 cm) were collected on the east side of Admiralty Bay, near Point Hennequin, using a Van-Veen Grab Sampler. Approximately 200 g of sediments of each sample were placed into Whirl-Pak bags. Superficial soil samples (0 - 5 cm) were collected on the proximities of EACF and then placed into Whirl-Pak bags (~200 g). Samples from Deception Island comprised surface sediments (0 - 5 cm) in an intertidal region near active fumaroles, with temperatures of 110 °C for FBA1, FBA2 and FBA3, and 112 °C for FBB1, FBB2 and FBB3. Fumarole sediments were placed into Whirl-Pak bags (~200 g).

The Continental Antarctica sampling site is located at West Antarctica, 250 km from the southwest border of the Ronne ice shelf and 670 km from the geographic South Pole, where the Brazilian module Criosfera 1 is located (S 84°00’, W 079°30’). Snow/firn samples were collected in an aseptic excavated pit structure near the Brazilian module. Six depths were collected between the surface and 200 cm, including 0 - 40 cm (C1), 40 - 85 cm (C2), 85 - 110 cm (C3), 110 - 160 cm (Crio4), 160 - 182 cm (Crio5), 182 - 200 cm (C6). Approximately 3 L of water of each sample were filtered in the Criosfera 1 module using a vacuum pump and 0.22 μm-membrane filters.

All samples collected in this study were immediately frozen at −20°C for molecular analysis. The description of environmental samples, the coordinates and sampling year are detailed in Supplementary Table 1.

### 2.2. DNA extraction and sequencing of the 16S rRNA gene

The 0.22 μm-membrane filters of seawater and snow samples were submitted to DNA extraction using DNeasy PowerWater Kit (Qiagen, Hilden, Germany). For sediment and soil samples, approximately 500 mg were submitted to DNA extraction using DNeasy PowerSoil Kit (Qiagen, Hilden, Germany). Approximately 10 g of fumarole sediments were submitted to DNA extraction using DNeasy PowerMax Soil Kit (Qiagen, Hilden, Germany). All extractions were performed according to the manufacturer’s instructions. Extracted DNA was quantified using Qubit dsDNA HS Assay (Thermo-Fisher Scientific, Waltham, U.S.A.) and Qubit Fluorometer 1.0 (Thermo-Fisher Scientific, Waltham, U.S.A.).

Total extracted DNA were sequenced using Illumina Miseq paired-end system 2 x 300 bp, with the primers 515F (5’-GTGYCAGCMGCCGCGGTAA-3’) and 806R (5’-GGACTACNVGGGTWTCTAAT-3’) (Caporaso et al., 2012) for fumarole sediment and snow samples, targeting the V4 region of the 16S rRNA gene, and the primers 515F (5’-GTGYCAGCMGCCGCGGTAA-3’) and 926R (5’-CCGYCAATTYMTTTRAGTTT-3’) (Quince et al., 2011) for seawater, soil and marine sediment samples, targeting the V4 and V5 regions of the 16S rRNA gene. Details of pairs of primers used for each sample are in Supplementary Table 1. Library construction and sequencing were performed by MR DNA (Molecular Research LP, Shallowater, TX, EUA). The library sequencing followed the Earth Microbiome Project protocol (Thompson et al., 2017).

### 2.3. Bioinformatics and statistical analyses

Reads were initially imported into the Quantitative Insights Into Microbial Ecology 2 software (Qiime2) (v.2020.2, https://docs.qiime2.org/) (Bolyen et al., 2019) and then evaluated according to quality. To be consistent among the different sequence datasets and pairs of primers used in our study, only forward sequences (R1) were processed, comprising the V4 region of the 16S rRNA gene. Based on the quality scores, the forward reads were truncated at position 230, and trimmed at the position 25 to remove the primer, using the q2-dada2-denoise script. DADA2 software was used to obtain a set of observed amplicon sequence variants (ASVs) (Callahan et al., 2017). Taxonomic classification was performed through feature-classifier classify-sklearn using the Silva v.138 database (Quast et al., 2013; Yilmaz et al., 2014). The alignment was performed by MAFFT v.7 (Katoh et al., 2002), using default parameters and the phylogenetic tree was built by FastTree (Price et al., 2009).

The Qiime2 output qza files were imported on R version 4.0.4 (R CORE TEAM) using the qiime2R package (https://github.com/jbisanz/qiime2R). Alpha and beta diversity metrics were computed through the phyloseq package (McMurdie and Holmes, 2012) on R at a rarefied sampling depth of 11,604 sequences. Statistical differences in alpha diversity indices were calculated by comparing sample types and location using the ANOVA test in stats package on R. Beta diversity was measured by weighted Unifrac distance and visualized via NMDS (non-metric multidimensional scaling) using the phyloseq package in R (version 3.6.3). Differences in the microbial community structure among sample types and location were tested by performing a permutational multivariate analysis of variance (PERMANOVA) on the community matrix (Anderson, 2001).

To observe the unique and shared ASVs by each sample type, the taxa abundance table was transformed to presence/absence. The number of shared ASVs by sample types was visualized using an UpSet plot, UpSetR package (Conway and Gehlenborg, 2019). The core microbiome of each sample type was considered as the shared ASVs within the sample type, which was visualized at order level through pie charts. The statistical package IndicSpecies (Cáceres et al., 2020) was used on R to identify microbial families whose abundance was significantly associated with a sample type.

Sequencing data were deposited in the National Center for Biotechnology Information Sequence Read Archives (SRA) under BioProject IDs XXXXX (IDs will be provided immediately after manuscript acceptance).

## 3. Results

### 3.1. Richness and alpha diversity

We obtained 4,781,877 valid sequences distributed among 5 sample types (habitats), including 3 samples of marine sediment, 6 samples of fumarole sediments, 6 samples of snow/firn, 52 samples of seawater and 27 samples of soil, totalizing 94 samples. A mean of 336 ASVs (SD ± 212) were detected for each sample. The values of ASVs, richness (Chao1) and alpha diversity (Shannon and InvSimpson) were statistically different (*p* <0.05) according to sample type, and not by location (*p* = 0.96 for Chao1, *p* = 0.44 for Simpson and *p* = 0.28 for InvSimpson). Richness and alpha diversity results are represented in Figure 2 and detailed in Supplementary Table 1.

**Figure 1.**
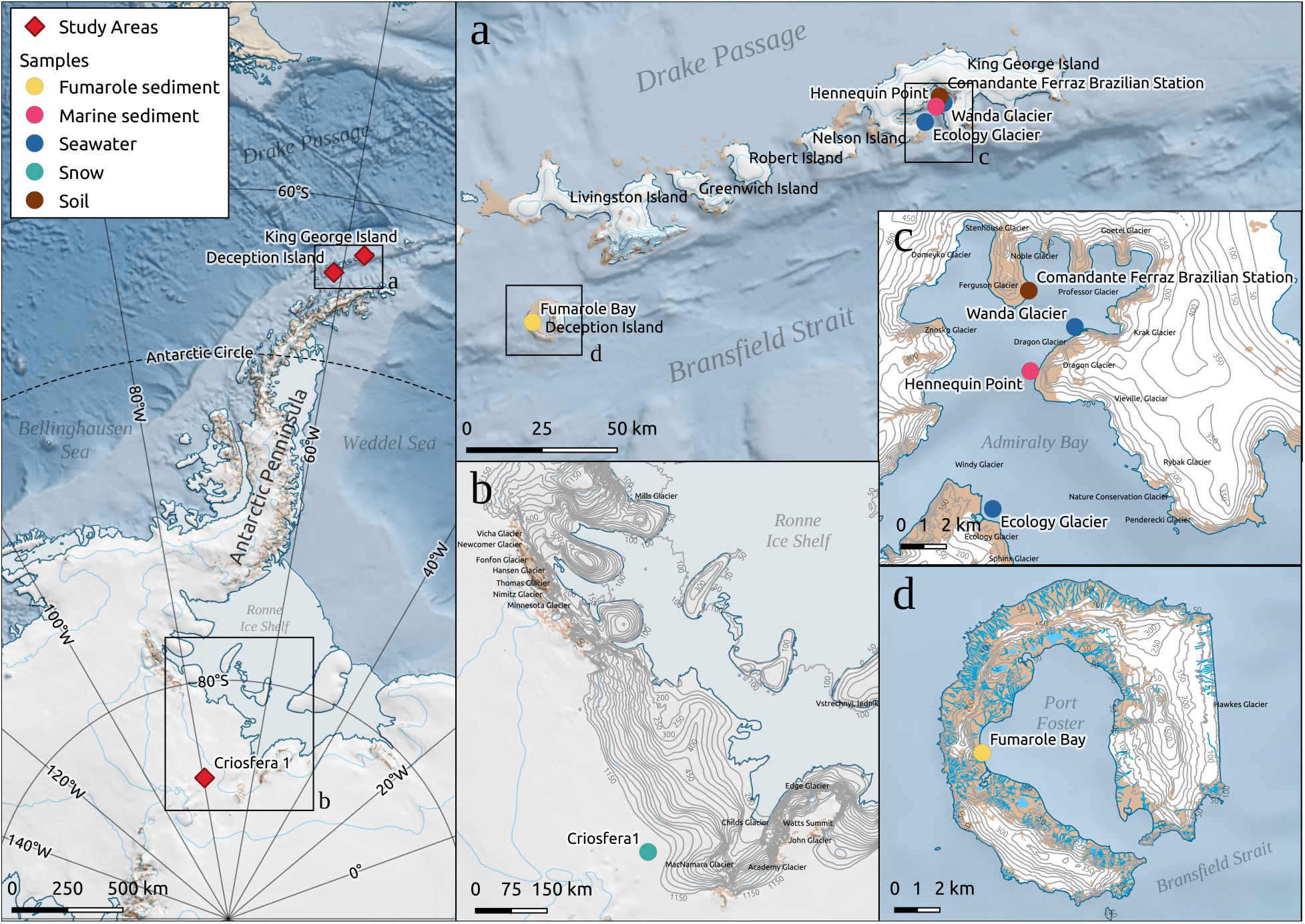
Study locations and sampling sites in the northwest region of Antarctica. The subfigures a, b, c and d represent, respectively, the South Shetland Islands region, the southwest border of the Ronne Ice Shelf, the Admiralty Bay in King George Island and the Deception Island. The red diamonds on the left side represent the three distinct study areas, and the circle represents the sample types by colors (yellow = fumarole sediment, pink = marine sediment, dark blue = seawater, light blue = snow, brown = soil). The map was made by using the Qgis software (QGIS.org 2021) and the Quantarctica data set (Matsuoka et al. 2018).

**Figure 2.**
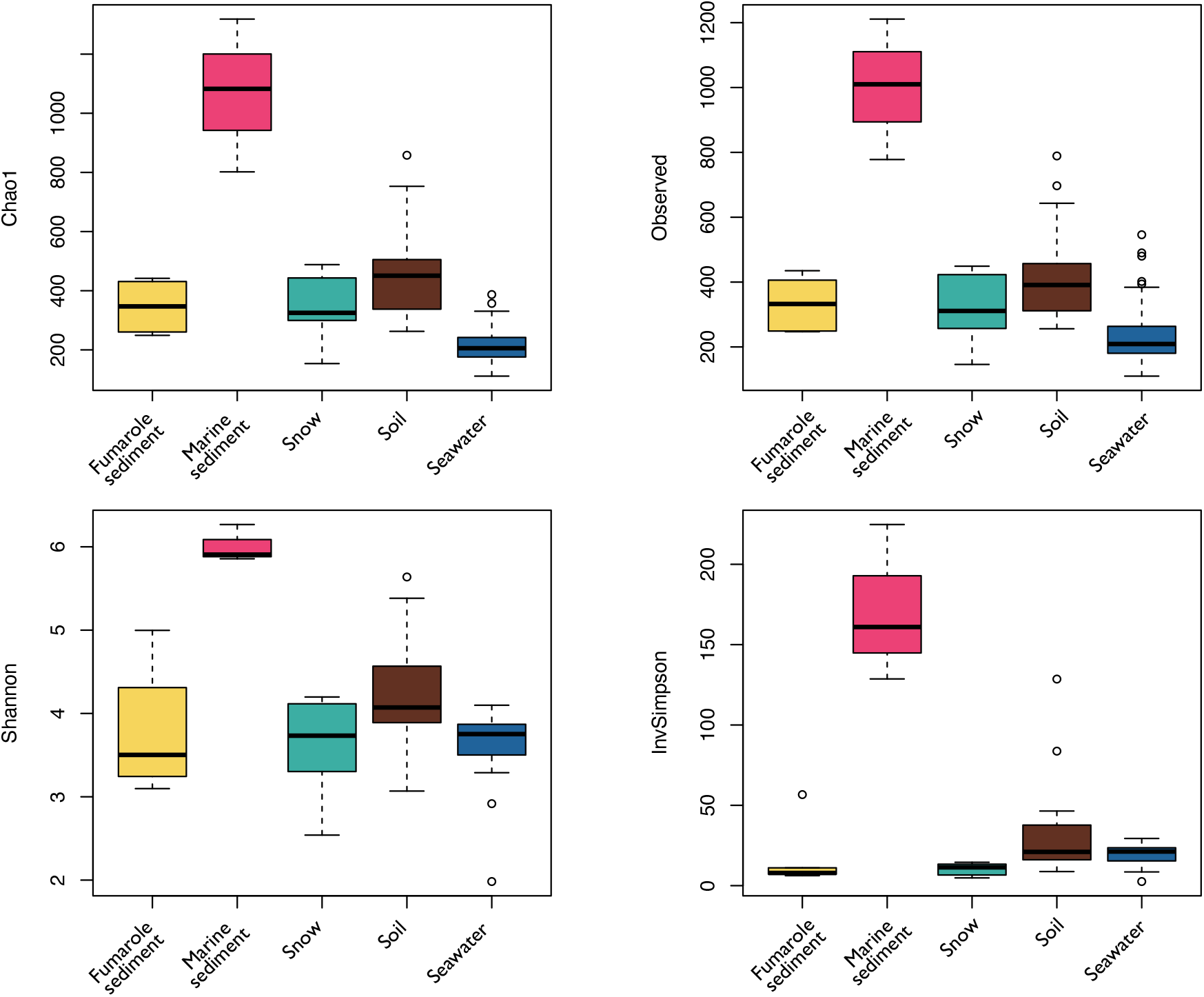
Alpha diversity analyses, including the number of ASVs (observed), the richness index of Chao1, and the alpha diversity indices of Shannon and InviSimpson. Samples are grouped by each habitat (sample type).

When grouped by location, the richness and alpha diversity values for the Antarctic continent samples were 333.32±116.73[SD] (Chao1), 3.60±0.63 (Shannon) and 10.30±3.78 (InvSimpson); 346.84±84.92 (Chao1), 3.78±0.75 (Shannon) and 16.29±20.29 (InvSimpson) for Deception Island; 323.72±213.57 (Chao1), 3.93±0.63 (Shannon) and 28.84±33.58 (InvSimpson) for King George Island.

When grouped by sample types, marine sediment samples exhibited the highest values of richness (Chao1= 1095.24±276.30[SD]) and alpha diversity (Shannon= 6.01±0.23; InvSimpson= 174.55±53.05), followed by soil samples (Chao1= 453.68±149.46; Shannon= 4.21±0.57; InvSimpson= 30.16±26.14). Fumarole sediments (Chao1= 349.38±84.70; Shannon= 3.79±0.74; InvSimpson= 16.46±20.48), snow (Chao1= 339.02±121.31; Shannon= 3.60±0.64; InvSimpson= 10.32±3.87) and seawater (Chao1= 210.77±44.89; Shannon= 3.66±0.33; InvSimpson= 19.83±5.18) exhibited the lowest values of richness and alpha diversity indices.

### 3.2. Beta diversity

Samples were clustered according to sample type and location through the weighted Unifrac distance analysis observed in NMDS (Figure 3). Seawater samples were grouped nearest from each other, as well as marine sediments. Samples of soil, fumarole sediment and snow exhibited a clustering pattern more distant from each other. Based on the PERMANOVA, samples were significantly influenced more by sample type (p<0.01, R^2^=0.61) than by location (p<0.01, R^2^=0.17).

**Figure 3.**
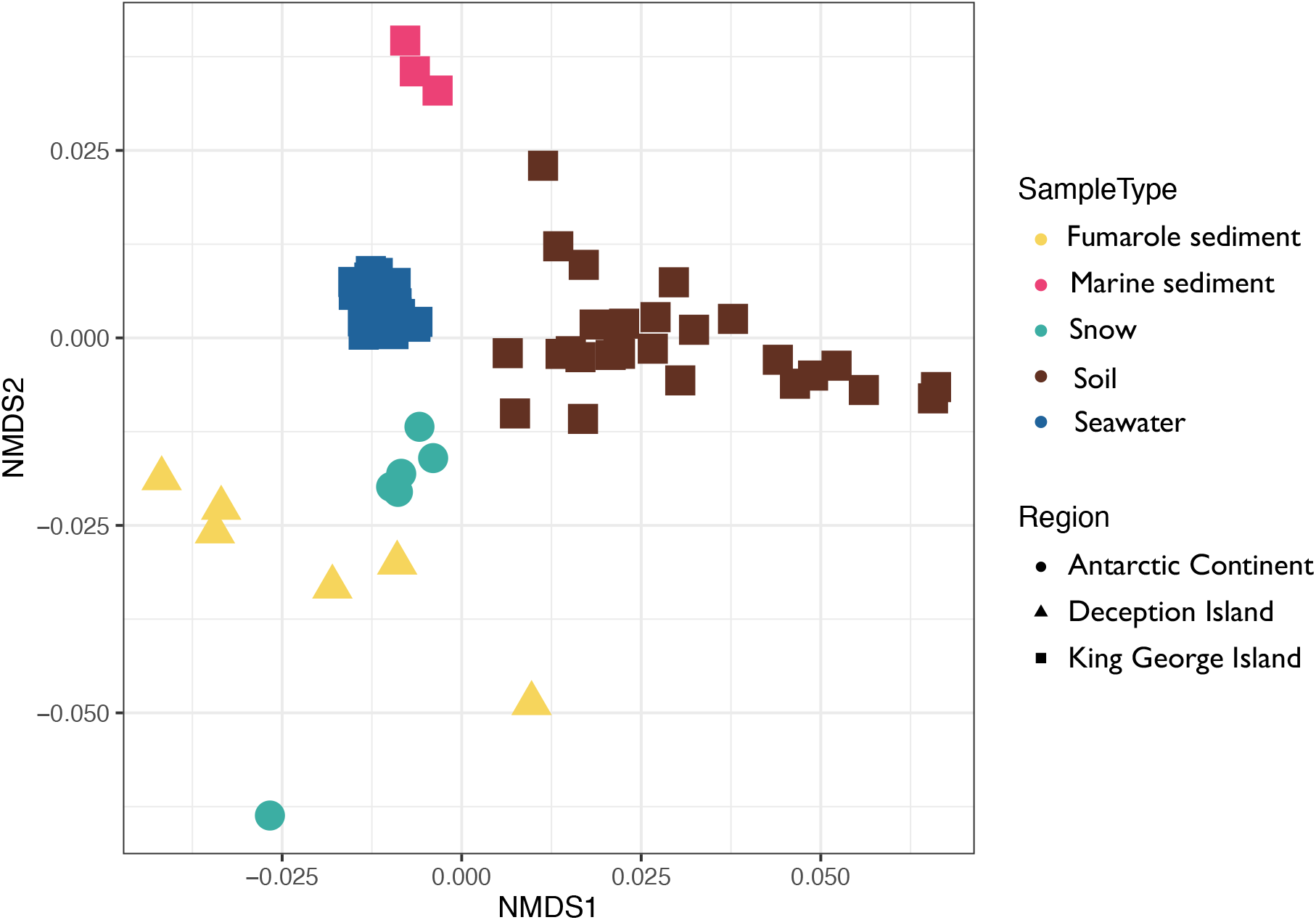
Non-metric multidimensional scaling (nMDS) ordination based on weighted UNIFRAC distances. The shapes represent the three main regions in Antarctica and colors the Antarctic habitats (sample types). Stress value=0.118.

### 3.3. Microbial community composition at phylum level

A total of 29 phyla were classified as abundant (> 1% of relative abundance) among our samples (Figure 4). The most abundant phyla in marine sediments were Proteobacteria (21.8±1.8%[SD]), Bacteroidota (19.9±5.0%), Acidobacteriota (14.0±2.9%), Verrucomicrobiota (11.8±2.4%), Actinobacteriota (9.3±1.1%), Chloroflexi (8.4±2.5%), Planctomycetota (4.1±1.4%), Gemmatimonadota (3.3±1.0%), Nitrospirota (2.2±0.8%) and Crenarchaeota (1.0±0.5%). In fumarole sediments, abundant phyla were classified as Aquificota (21.6±11.5%), Proteobacteria (21.1±13.6%), Crenarchaeota (13.6±9.5%), Firmicutes (11.3±8.2%), Deinococcota (6.0±5.9%), Actinobacteriota (3.8±1.4%), Patescibacteria (0.02±1.2%), Bacteroidota (1.7±1.0%), Chloroflexi (1.6±1.8%), Verrucomicrobiota (1.5±0.5%) and Nanoarchaeota (1.1±0.7%). The most abundant phyla in snow samples were Proteobacteria (77.6±17.5%), followed by Actinobacteriota (9.0±14.6%), Firmicutes (7.5±3.9%) and Bacteroidota (1.5±0.8%). For water samples, only two phyla were abundant: Proteobacteria (62.8±5.8%) and Bacteroidota (35.4±5.8%). Abundant phyla in soil samples were Proteobacteria (63.2±0.9%), Bacteroidota (22.7±7.7%), Actinobacteriota (7.2±3.4%) and Acidobacteriota (3.0±3.6%).

**Figure 4.**
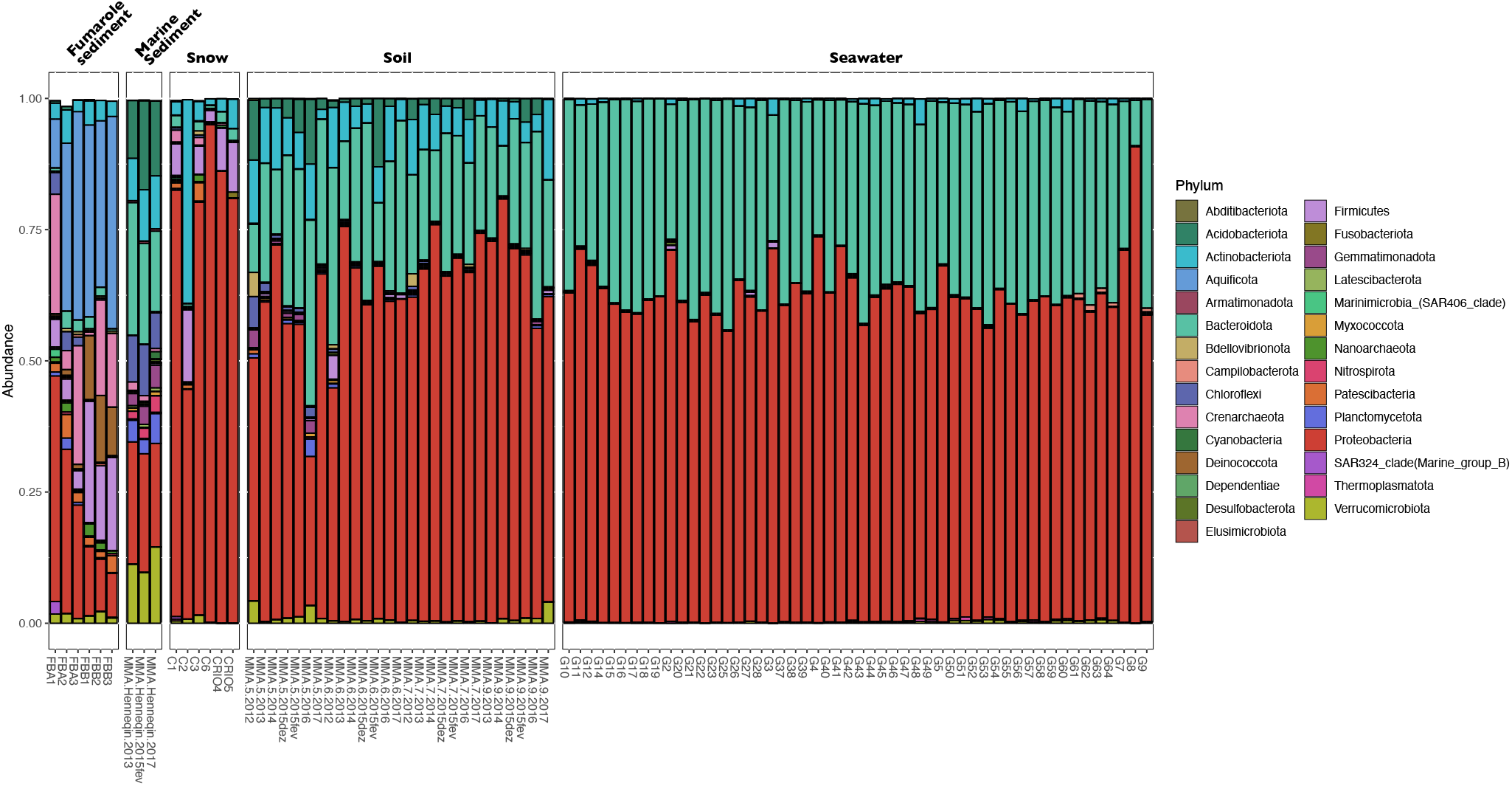
Microbial community composition grouped by each Antarctic habitat (sample type). The figure shows the relative abundance of bacterial and archaeal taxonomic groups at phylum level. Only phylum with more than 0.1% of abundance are represented. Sequences were taxonomically classified using the Silva database v. 138.

### 3.4. Shared ASVs and core microbiome

The number of shared ASVs among sample types are represented in the upset plot of Figure 5. In general, communities from snow shared more ASVs with fumarole sediments (157 ASVs) and seawater (48 ASVs), whereas soil communities shared more ASVs with marine sediments (378 ASVs) and seawater (115 ASVs). The pie charts (Figure 5) represent the taxonomic classification of ASVs (at order level) that were considered the core microbiome of each sample type. The core microbiome indicates the microbial taxa that are particularly widespread within a sample group. The results of core microbiome per sample type are detailed in Supplementary Table 2.

**Figure 5.**
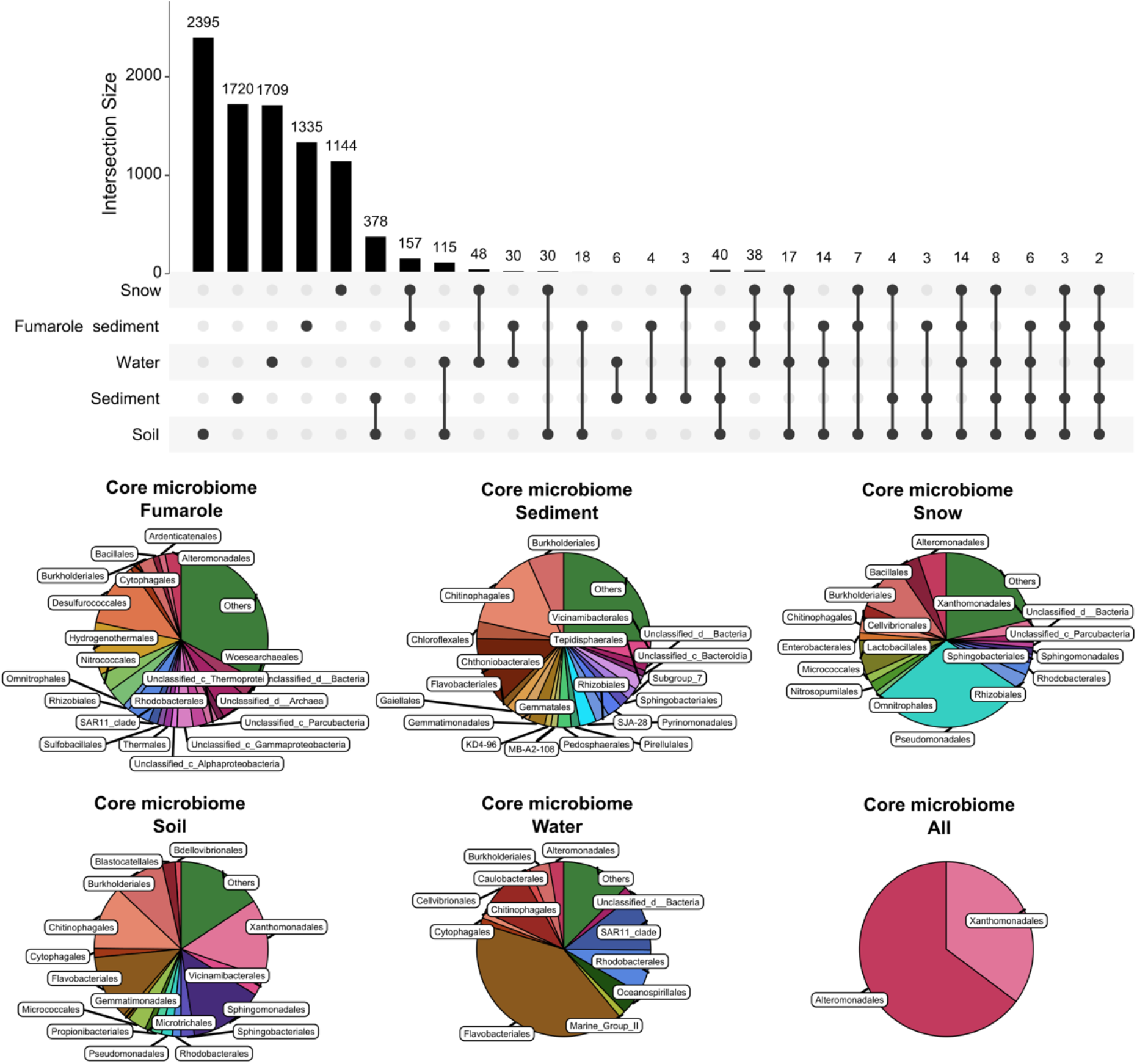
Upset plot composed by ASVs identified among sample types. Circles indicate sample types. Black lines connecting circles indicate shared ASVs. Vertical bars indicate intersection size (number of ASVs) on each set. Pie charts show microbial composition specific to each sample type (orders with abundance > 1%) and those shared among all sample types or habitats (core microbiome).

The core microbiome of marine sediments was composed mainly by the orders Chitinophagales (14.8%), Chthoniobacterales (12.5%), Burkholderiales (6.6%), Vicinamibacterales (3.7%), Chloroflexales (3.2%), Pyrinomonadales (3.0%), Gemmatimonadales (3.0%), among others. For fumarole sediments, the core microbiome was composed by orders such as Desulfurococcales (11.8%), Hydrogenothermales (9.6%), Unclassified_Bacteria (5.4%), Rhodobacterales (4.3%), Woesearchaeales (3.7%), Omnitrophales (3.5%), Nitrococcales (3.4%), among others. The core microbiome of snow samples included Pseudomonadales (29.5%), Burkholderiales (9.0%), Lactobacillales (6.4%), Alteromonadales (5.2%), Bacillales (4.2%), Chitinophagales (2.9%), among others. Seawater samples exhibited as the core microbiome the orders Flavobacteriales (40.7%), SAR11_clade (10.6%), Cellvibrionales (9.9%), Rhodobacterales (8.1%), Oceanospirillales (4.3%), Burkholderiales (3.7%), Alteromonadales (2.7%), Marine_Group_II (1.2%), among others. The core microbiome of soil samples comprised the orders Xanthomonadales (14.4%), Sphingomonadales (13.8%), Flavobacteriales (12.0%), Chitinophagales (11.8%), Burkholderiales (9.4%), Vicinamibacterales (3.5%), among others. Finally, the core microbiome when considered all samples was composed by two orders: Xanthomonadales (35.1%) and Alteromonadales (64.8%).

### 3.5. Microbial indicators for each sample type

By using the R package IndicSpecies we were able to identify the families significantly associated with each sample type, which are represented in Figure 6 and detailed in Supplementary Table 3. Marine sediments was the sample type which exhibited the highest number of indicators, totalizing 81 families classified within 22 phyla, such as Anaerolineaceae (Chloroflexi), Pyrinomonadaceae (Acidobacteriota), Holosporaceae (Proteobacteria) and Gaiellaceae (Actinobacteria). A total of 12 families were indicators for fumarole sediments: lineage_IV within Elusimicrobiota, Pyrodictiaceae, Hydrogenothermaceae, Candidatus_Zambryskibacteria, Desulfurococcaceae, Candidatus_Nomurabacteria, Acidilobaceae, SAR202_clade, Methylomirabilaceae, Thermaceae, Thermicanaceae and Woesearchaeales. For snow samples, 4 families were considered as indicators, classified as Oleiphilaceae (Proteobacteria), Burkholderiaceae (Proteobacteria), Bifidobacteriaceae (Actinobacteriota) and Exiguobacteraceae (Firmicutes). Eleven families were indicators of seawater samples, which were classified as Parvibaculaceae, OCS116_clade, Cryomorphaceae, OM182_clade, NS7_marine_group, Clade_III (SAR11_clade), Marine_Group_II, Psychromonadaceae, Arcobacteraceae, SAR116_clade and uncultured family. Finally, 4 families were indicators of soil samples, which belonged to NRL2 (Proteobacteria), Demequinaceae (Actinobacteriota), Iamiaceae (Actinobacteriota) and Immundisolibacteraceae (Proteobacteria).

**Figure 6.**
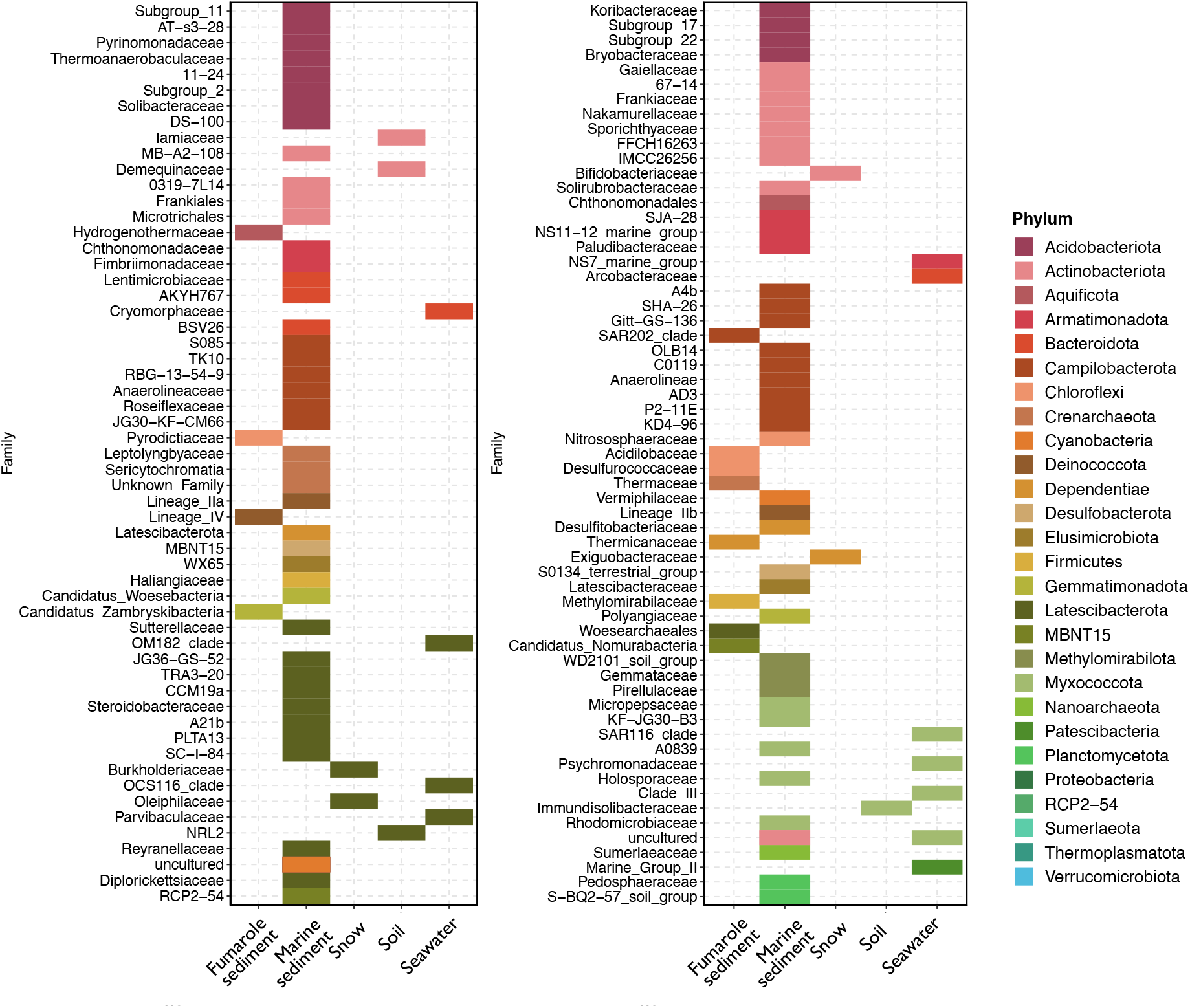
Indicator families identified as significantly associated with each sample type (habitat), calculated using the R package IndicSpecies. The colors represent the phyla classifications of each family.

## 4. Discussion

In our study, we were able to describe the core microbiome and the microbial indicators of five Antarctic habitats located at both Maritime and Continental Antarctica. Our results showed that marine sediment was the habitat which harbored the highest microbial diversity. We observed a significant difference of microbial community structure according to each habitat, showing that despite geographical distances, the environmental conditions act as strong pressures for selecting specific microbial taxa.

### 4.1. Microbiome of marine sediments from King George Island

Globally, marine sediments cover 70% of Earth’s surface and are thought to be a larger biomass reservoir than seawater, counting for 0.18 to 3.6% of the total living biomass of the Earth (Kallmeyer et al., 2012; Parkes et al., 2014). In Antarctica, the estimation of the microbial biomass in marine sediments is still poorly understood. The microbial abundance in marine sediments is frequently associated with depth patterns, generally decreasing with increasing depth. A recent study estimated a bacterial and archaeal richness in marine sediment between 4.03 × 10^4^ to 3.30 × 10^6^ ASVs (Hoshino et al., 2020). These values were comparable to the richness estimated for topsoil and seawater samples, which comprised 7.88 × 10^4^ to 1.69 × 10^7^, and 3.00 × 10^4^ to 1.69 × 10^6^, respectively (Hoshino et al. 2020). In the present study, marine sediments showed the highest microbial richness (1.09 × 10^3^ ASVs) when compared to the other habitats, but exhibited lower values than those estimated by Hoshino et al. (2020). It is plausible to observe these contrasts between the global estimations for marine sediments and our richness results, since benthic communities in Antarctica have to adapt to the environmental extreme conditions, such as the prevalent low temperatures, freeze and thaw cycles, low nutrient input, and high salinity (Bölter et al., 2002; Convey et al., 2009). These conditions produce narrow microbial niches and demand specific adaptive mechanisms for microbial growth and survival (Cowan et al., 2014).

Also, this pattern probably reflects the more stable temperatures in the sediments (when compared to other Antarctic habitats), where communities should fluctuate little seasonally, and then more microbial taxa could be able to survive. Other possibilities to explain the highest diversity in marine sediments could be due to the contribution of the communities from soil and snow, which reach inlet waters as results of glacier defrost, or due to cell deposition by descendant of pelagic communities, which could be buried and preserved for long periods (Hoshino et al., 2020).

The core microbiome of marine sediments from Admiralty Bay (King George Island) had the prevalence of Bacteroidota, Verrucomicrobiota, Acidobacteria, Chloroflexi, Gemmatimonadota and Proteobacteria, in which some members of these phyla have been previously described in marine sediments of the Antarctic Peninsula (Foong et al., 2010; Li et al., 2020; Powell et al., 2003). Franco et al. (2017) revealed a high prevalence of heterotrophic gammaproteobacterial phylotypes in the marine sediments of Admiralty Bay, but also reported the presence of taxa from Bacteroidota, Verrucomicrobiota, Acidobacteria, Chloroflexi, Gemmatimonadota phyla.

Among the microbial families observed as indicators of marine sediments, Anaerolineaceae (Chroloflexi) have been previously described as abundant in marine sediments, being involved with hydrocarbon degradation (Fincker et al., 2020). In addition, we also observed Pyrinomonadaceae as an indicator of marine sediments, which members were previously observed in diesel contaminated soil samples from King George Island (Gran-Scheuch et al., 2020), and also in other extreme environments, such as semi-arid savannah and volcanic soils (Pascual et al., 2018). This bacterial family comprises aerobic and chemoheterotrophic mesophiles or thermophiles, capable of growing in mildly acidophilic environments (Dedysh and Damsté, 2018).

### 4.2. Microbiome of fumarole sediments from Deception Island

Previous studies have indicated that temperature is one of the major drivers of microbial communities’ diversity and structure (e.g. Price and Giovannelli, 2017; Sharp et al., 2014). Geothermal and hydrothermal ecosystems have been considered as “open-air” laboratories for revealing the responses of microbial communities to temperature gradients (e.g. Antranikian et al., 2017; Bendia et al., 2018; Ward et al., 2017). One of the most interesting ecosystems to explore temperature-adapted extremophiles (psychrophiles, thermophiles, and hyperthermophiles) are the polar volcanoes, where we can find extreme temperature and geochemical gradients over very short distances (Herbold et al., 2014). In Antarctica, a recent study showed that Deception Island volcano harbor different extremophilic lineages, which were strongly driven by steep temperature gradients (from 0 to 98 °C) (Bendia et al., 2018).

In our study, the fumarole sediments from Fumarole Bay on Deception Island, which comprised the temperatures of 110 °C and 112 °C, exhibited as the core microbiome mostly bacterial and archaeal lineages related to thermophiles and hyperthermophiles, such as those within the orders Hydrogenothermales, Sulfobacilalles, Desulfurococcales and Thermales. Further, the indicator families of fumarole sediments belong to thermophiles and hyperthermophiles (Pyrodictiaceae and Hydrogenothermaceae), and to spore-forming bacteria from Firmicutes phylum (Carnobacteriaceae). Pyrodictiaceae comprises members which are autotrophic anaerobes, hydrogen-oxidizers, denitrifiers and iron-reducers, whereas Hydrogenothermaceae are usually aerobes or anaerobes, autotrophs, sulfur-oxidizers and denitrifiers (Zeng et al., 2021). Our results indicate that, despite the geographic isolation and the predominantly cold habitats in Antarctica, the hyperthermophilic temperatures act as strong pressures on selecting hyperthermophilic lineages, which showed to be widespread across these fumaroles, as also observed by Bendia et al. (2018). By comparing Deception communities with continental geothermal systems in Antarctica, such as Tramway Ridge in Mount Erebus (Herbold et al., 2014; Soo et al., 2009), few taxa are shared, mainly related to Chloroflexi and Planctomycetes. Pyrodictiaceae and Hydrogenothermaceae lineages were found in geothermal systems, and in shallow and deep-sea hydrothermal vents, such as those in Mariana Volcanic Arc (Nakagawa et al., 2006), Manus Basin, New Guinea (Takai et al., 2001), Vulcano, Italy (Stetter et al., 1983), Tachibana Bay, Japan (Takai and Sako, 1999), and near Tonga subduction zone in the Southwestern Pacific (Ferrera et al., 2014).

### 4.3. Microbiome of snow from West Continental Antarctica

The continental snow represents a dynamic habitat where microorganisms encounter low temperatures, variability in surface UV radiation and limited water and nutrients availability (Larose et al., 2013). Previous studies, including from non-polar environments, identified Proteobacteria, Bacteroidetes, Firmicutes and Cyanobacteria as the dominant taxa in snow habitats in Antarctica (Antony et al., 2016; Lopatina et al., 2016; Malard et al., 2019; Michaud et al., 2014; Yan et al., 2012), Arctic (Harding et al., 2011; Hell et al., 2013; Larose et al., 2013; Maccario et al., 2014), Austria (Battin et al., 2001), Canada (Boyd et al., 2011) and Svalbard (Zarsky et al., 2013). Although previous studies investigated Antarctic snow, few have focused on microbial diversity and distribution, with these studies limited to specific locations leaving the vast majority of the continent unexplored (Boetius et al., 2015; Hodson et al., 2017; Luo et al., 2020).

In the present study, the snow from West Antarctica (near Brazilian module Criosfera1) exhibited as the core microbiome bacterial lineages related to Proteobacteria, especially Alphaproteobacteria and Gammaproteobacteria, and also several orders related to heterotrophs, such as Alteromonadales, Bacillales, Burkholderiales and Chitinophagales, which is in accordance with previous studies on the Antarctic snow microbial community (Michaud et al., 2014; Antony et al., 2016; Lopatina et al., 2016). We detected one archaeal taxa as the core microbiome in snow samples, assigned within the order Nitrosopumilales (Crenarchaeota), while a previous study (Antony et al., 2016) identified only Halobacteriaceae (Euryarchaeota) in snow samples from East Antarctica. The detection of Nitrosopumilales across a variety of temperature and saline gradients, suggests that its members have the ability to adapt to hot and cold habitats, as well as to terrestrial and marine ecosystems (Bendia et al., 2018; Learman et al., 2016; Lezcano et al., 2019; Pessi et al., 2015). The family indicators for snow samples were Oleiphilaceae, Burkholderiaceae, Bifidobacteriaceae and Exiguobacteraceae, whose members are often aerobes and heterotrophs (Biavati and Mattarelli, 2018; Coenye, 2014; Vishnivetskaya et al., 2009; Yakimov and Golyshin, 2014), and commonly present in soil habitats from Antarctica (Buelow et al., 2016; Chaturvedi et al., 2008; Pearce et al., 2012), except for Oleiphilaceae, which were predominantly found in deep marine sediments and are known to be hydrocarbon degraders (Bacosa et al., 2018; Golyshin et al., 2002).

It is still not clear if the presence of these bacteria and archaea in snow habitats reflects their ability to adapt and survive in extreme conditions (Edwards et al., 2014), or whether their high predominance in other Antarctic ecosystems favors their aeolian dispersion and preservation along surface habitats in the cryosphere (Archer et al., 2019). Previous studies suggested that soil microorganisms are the primary sources of snow microbial communities of the West Greenland Ice Sheet (Cameron et al., 2015) and Arctic (Cuthbertson et al., 2017; Šantl-Temkiv et al., 2018). Previous studies indicated the dominance of Proteobacteria and Firmicutes in airborne microbial communities in Antarctica (Bottos et al., 2014; Pearce et al., 2010), and the study by Malard et al., (2019) identified similarities between snow and airborne microbial communities in continental Antarctica, which suggests the importance of long-distance dispersal in seeding continental Antarctic snow ecosystems.

### 4.4. Microbiome of soils from King George Island

Although the ice-free areas comprise less than 0.3% of the total Antarctic area, soils are the most studied microbial habitat in Antarctica (Cowan et al., 2014). Soil habitats represent a wide variety of landforms and geochemistry, in which Proteobacteria and Actinobacteria showed to be dominant (Babalola et al., 2009; Makhalanyane et al., 2013). Archaeal taxa in Antarctic soils showed to be a negligible portion of the total microbial community and have likely a minimal role in soil processes (Cowan et al., 2014). A similar pattern was observed among our soil samples from King George Island, where Acidobacteriota, Actinobacteriota, Bacteroidota and Proteobacteria were the most abundant phyla, while several heterotrophic bacterial families, such as Pseudomonadales, Flavobacteriales, Cytophagales, Chitinophagales, comprised the core microbiome. Wang et al. (2015) also found the predominance of Proteobacteria, Actinobacteria, Acidobacteria, and Verrucomicrobia in four soil types at Fildes Region, King George Island, including pristine and human-impacted soils. Flavobacteriales members are widespread in terrestrial and marine Antarctic ecosystems, and the genus *Flavobacterium* have shown to play an important role in remineralization processes mainly due to its strong macromolecular hydrolytic capabilities (McCammon and Bowman, 2000). In contrast to our results, Ramos et al. (2019) showed a dominance of Firmicutes in soils from eleven regions of Admiralty Bay, King George Island. Differences in microbial composition of ecologically comparable soils from King George Island suggest a high level of spatial heterogeneity in prokaryotic diversity, as previously indicated by (Almela et al., 2021).

The indicator taxa of soil samples comprised four families, classified as Iamiaceae and Demequinaceae, both belonging to Actinobacteriota phylum and with members isolated from marine environments (Kurahashi et al., 2011; Ue et al., 2011), and NRL2 and Immundisolibacteraceae, which have lineages capable of hydrocarbon degradation (Corteselli et al., 2017). Since our soil samples were collected near Comandante Ferraz Brazilian Antarctic Station, the presence of hydrocarbon degraders might indicate an anthropogenic influence on microbial communities of the surrounding soil. Further, the presence of marine bacteria in soils from King George Island indicates that the ocean might be an important source of biological input to terrestrial environments, as suggested by Chong et al. (2012).

### 4.4. Microbiome of seawater from King George Island

Microbial communities along seawater samples from Admiralty Bay were very similar, even when comparing the superficial, intermediate and bottom depths. We observed as the core microbiome several marine orders, such as Alteromonadales, Oceanospirillales, SAR11 clade, Flavobacteriales, Rhodobacterales and the archaeal Marine Group II. These groups also showed to be abundant in shallow waters of the Bransfield Strait (Signori et al., 2018, 2014). Alteromonadales and Oceanospirillales are known to play an important role in organic carbon degradation by the production of extracellular hydrolytic enzymes (Dang et al., 2009). Some members of Oceanospirillales are also potential chemoautotrophs due to the presence of carbon fixation genes (Calvin Cycle pathway) (DeLorenzo et al., 2012). Although several members of the seawater community from Admiralty Bay were very similar to those found in surface waters of Bransfield Strait (Signori et al., 2018), we did not detect some key taxa, such as those within ammonia-oxidizing Archaea (Thaumarchaeota). Thaumarchaeota lineages were indeed detected in high abundance at surface colder waters of the Southern Ocean (~ −1 °C) (Signori et al., 2018), which might explain why they were not found in the warmer waters from Admiralty Bay. Further, the high number of Rhodobacterales members in our seawater samples might be explained because they are primary colonizers of particulate organic matter (Dang et al., 2009), which become more available by the processes of glaciers melting during summer.

Among the 11 families assigned as indicators of seawater samples, the majority include uncultivated marine lineages, such as OM182 clade, OCS1116 clade and NS7 marine group, whose metabolic capabilities and roles in biogeochemical cycles are still unknown. The archaeal Marine Group II was also assigned as an indicator of seawater and comprises uncultivated lineages generally more common in surface waters that are potentially phototrophs due to the presence of proteorhodopsin genes (Pereira et al., 2019). Further, several members of the seawater microbiome have shown to contribute to important ecological processes in oligotrophic and cold waters, such as to biomass accumulation and to remineralization of organic matter, so that any environmental changes could strongly affect their functioning in biogeochemical cycles (Tonelli et al., 2021), with possible cascading effects on higher trophic levels (Signori et al., 2018).

## 5. Conclusion

In conclusion, our study showed that in Antarctica, the microbiome of each terrestrial and marine habitats here analysed, harbors specific bacterial and archaeal taxa. In fumarole sediments, we found the higher proportion of archaeal taxa, which were mostly related to hyperthermophiles, while in soil samples archaeal lineages were very low abundant or absent. Marine sediments showed the highest microbial diversity and then more taxa indicators when compared to the other habitats. Surprisingly, although geographically distant, the continental snow samples exhibited common taxa in comparison to the habitats from the Antarctic Peninsula, which suggests long-distance dispersal processes occurring from the Peninsula to the Continent. Seawater communities showed to harbor similar taxa from those previously described for Bransfield Strait, with the absence of some taxa, such as ammonia-oxidizing Thaumarchaeota members. The description and proposal of key taxa from different Antarctic microbiomes are important for further studies aiming to elucidate which environmental factors drive those microbial communities, as well as to give insights about the interplay of microbial assemblages among the Antarctic ecosystems.

## Acknowledgements

We thank the captain and the crew of the research polar vessel Almirante Maximiano, and the chief and team of the Comandante Ferraz Brazilian Antarctic Station for their support in sampling during the OPERANTARs XXX to XXXV. We thank the Criosfera 1 team, in special Dr. Emanuele Kuhn, for their support in sampling during OPERANTAR XXXII. We are very thankful to LECOM’s research team, and Rosa C. Gamba for their scientific support.

## Supplementary Table Legends

Supplementary Table 1. Detailed description of environmental samples, including Sample IDs, location, coordinates, sample types, depth, sampling year, name of the project, DNA extraction method, pairs of primers, and diversity indices assigned for each sample.

Supplementary Table 2. Results from core microbiome analysis, at order level, represented as percentages (%) by each sample type (habitat).

Supplementary Table 3. Results from IndicSpecies analysis, at family level, including the number of p values for each taxa and grouped by sample type (habitat).

